# In vivo targeted gene delivery using Adenovirus-antibody molecular glue conjugates

**DOI:** 10.1101/2025.01.31.635969

**Authors:** Paul J. Rice-Boucher, Elena A. Kashentseva, Igor P. Dmitriev, Hongjie Guo, Jacqueline M. Tremblay, Charles B. Shoemaker, David T. Curiel, Zhi Hong Lu

**Affiliations:** Department of Radiation Oncology, Washington University School of Medicine, St. Louis, MO, USA; Department of Biomedical Engineering, McKelvey School of Engineering, Washington University in Saint Louis, St. Louis, MO, USA; Tiger Biologics LLC, St. Louis, MO, USA; Department of Infectious Disease and Global Health, Tufts Cummings School of Veterinary Medicine, North Grafton, MA, USA

**Keywords:** Gene delivery, adenovirus, molecular glue, SpyTag, SpyCatcher, DogTag, DogCatcher

## Abstract

Safe and efficient nucleic acid delivery to targeted cell populations remains a significant unmet need in the fields of cell and gene therapy. Towards this end, we pursued Adenoviral vectors genetically modified with the “DogTag” molecular glue peptide, which forms a spontaneous covalent bond with its partner protein, “DogCatcher”. Genetic fusion of DogCatcher to single-domain or single-chain antibodies allowed covalent tethering of the antibody at defined locales on the vector capsid. This modification allowed simple, effective and exclusive targeting of the vector to cells bound by the linked antibody. This dramatically enhanced gene transfer into primary B and T cells *in vitro* and *in vivo* in mice. These studies form the basis of a novel method for targeting Adenovirus that is functional in stringent *in vivo* contexts and can be combined with additional well characterized Adenovirus modifications towards applications in cell engineering, gene therapy, vaccines, oncolytics, and others.

## Introduction

Effective genetic medicine requires the safe and efficient delivery of nucleic acids to cells and tissues of interest. Decades of research has yielded numerous vehicles for delivery of both DNA and RNA, including non-viral liposomes, lipid nanoparticles and others, and viral vectors such as adeno-associated viruses (AAVs), Adenoviruses (Ads), and lentiviruses^1–3^. Despite this, a vector capable of efficient and targeted delivery of genes to cells of interest remains elusive. This requirement is of particular importance to support the deployment of gene editors *in vivo*, as expression of the editing machinery in off-target regions may lead to undesirable side-effects. Despite these challenges, direct *in vivo* gene delivery remains an attractive option, potentially circumventing the costly and complicated procedures required for *ex vivo* cell engineering and enabling novel therapies deliverable to underserved patient populations^4–7^. Recognition of this has led to extensive research into *in vivo* T cell engineering, and a single report describing *in vivo* B cell engineering^8–15^. Targeted gene transfer into these cell types will thus likely be of strong utility.

Our group and others have undertaken extensive research into Adenoviral vectors as nucleic acid delivery vehicles. Ads are non-enveloped double-stranded DNA viruses generally responsible for the common cold and are amongst the best-studied DNA vectors. Ads have a long track-record in the clinic and have been successfully deployed for cancer immunotherapy and vaccines against infectious disease, with estimated doses delivered for COVID-19 ranging in the hundreds of millions^16,17^. Ads thus represent a potentially low-cost and safe vector for *in vivo* cell and gene therapy. Numerous manuscripts have been published on the targeting of Ads both *in vitro* and *in vivo*, but these approaches generally rely on the use of small peptides, adaptors, complicated genetic engineering, or the use of bespoke reagents such as DARPINs or single-domain antibodies (sdAbs)^18–20^. The ideal delivery vehicle would embody a single-component vector capable of being grown to high titers and targeted with easily obtained reagents.

With these goals in mind, we pursued the development of Ad vectors modified with SpyCatcher/SpyTag family molecular glues. We demonstrate the production of a Human Adenovirus serotype C5 (Ad5) vector with the “DogTag” peptide genetically incorporated in the fiber protein. Through this moiety we covalently attach various antibody species fused with DogCatcher to the virus surface and show that this linkage results in vector retargeting both *in vitro* and *in vivo* in primary B and T cells.

## Results

### Development of molecular glue driven antibody conjugation to the virus capsid

SpyCatcher/SpyTag protein-peptide partners spontaneously form a covalent bond under physiological conditions and have been used in numerous protein engineering studies^21,22^. Our group previously demonstrated that Simian Adenovirus serotype 36 (SAd36) could be derivatized with SpyTag at each of the major capsid proteins, including fiber, hexon, and pIX – the fiber protein is responsible for the initial binding of the virus to cells, while the hexon is the main structural protein forming the capsid. pIX interlaces the hexon and stabilizes the overall structure^18^. These derivatives could be linked with a Cas9-SpyCatcher fusion protein to achieve gene editing *in vitro*^23^. We thus hypothesized that a similar approach might be useful for targeting – by linking an antibody on the virus capsid through SpyCatcher/SpyTag chemistry, we might be able drive viral uptake through binding of the antibody to the appropriate receptor on a cell surface.

We selected B cells as a first choice for targeting due to our previous work on this cell type and standing interest in achieving *in vivo* engineering for control of infectious diseases^24^. We initially attempted targeting through our existing SpyTag modified SAd36 vectors using an sdAb targeting murine CD40 fused with SpyCatcher (F8SpC), but found these vectors were completely unable to access the immunocyte population in pilot *in vivo* studies and were thus likely inappropriate for our ultimate aims. Furthermore, concurrent with this work a study describing Ads engineered with the loop-friendly molecular glue “DogTag” was published^25,26^. Based on our previous positive results with Ad5 based vectors for *in vivo* gene delivery to B cells, we thus decided to design an Ad5 vector incorporating DogTag in the Ad5 fiber protein (Ad5FDgT). We selected the HI loop within the fiber for DogTag insertion, as this site has previously been used for insertion of peptides (**Fig. 1a** and **1b**)^27,28^. We found that this vector was easily upscaled in standard HEK293 cells and yielded titers comparable to the isogenic unmodified Ad5 vector. SDS-PAGE analysis revealed identical protein band patterns between Ad5 and Ad5FDgT, with the exception of the expected band shift from the insertion of DogTag in the fiber (**Fig. 1d**). We note that the wild-type Ad5 fiber and pIIIa proteins overlap just below the 70kDa protein ladder marker, whereas the DogTag modified fiber separates from the pIIIa protein due to its larger molecular weight.

**Figure 1:**
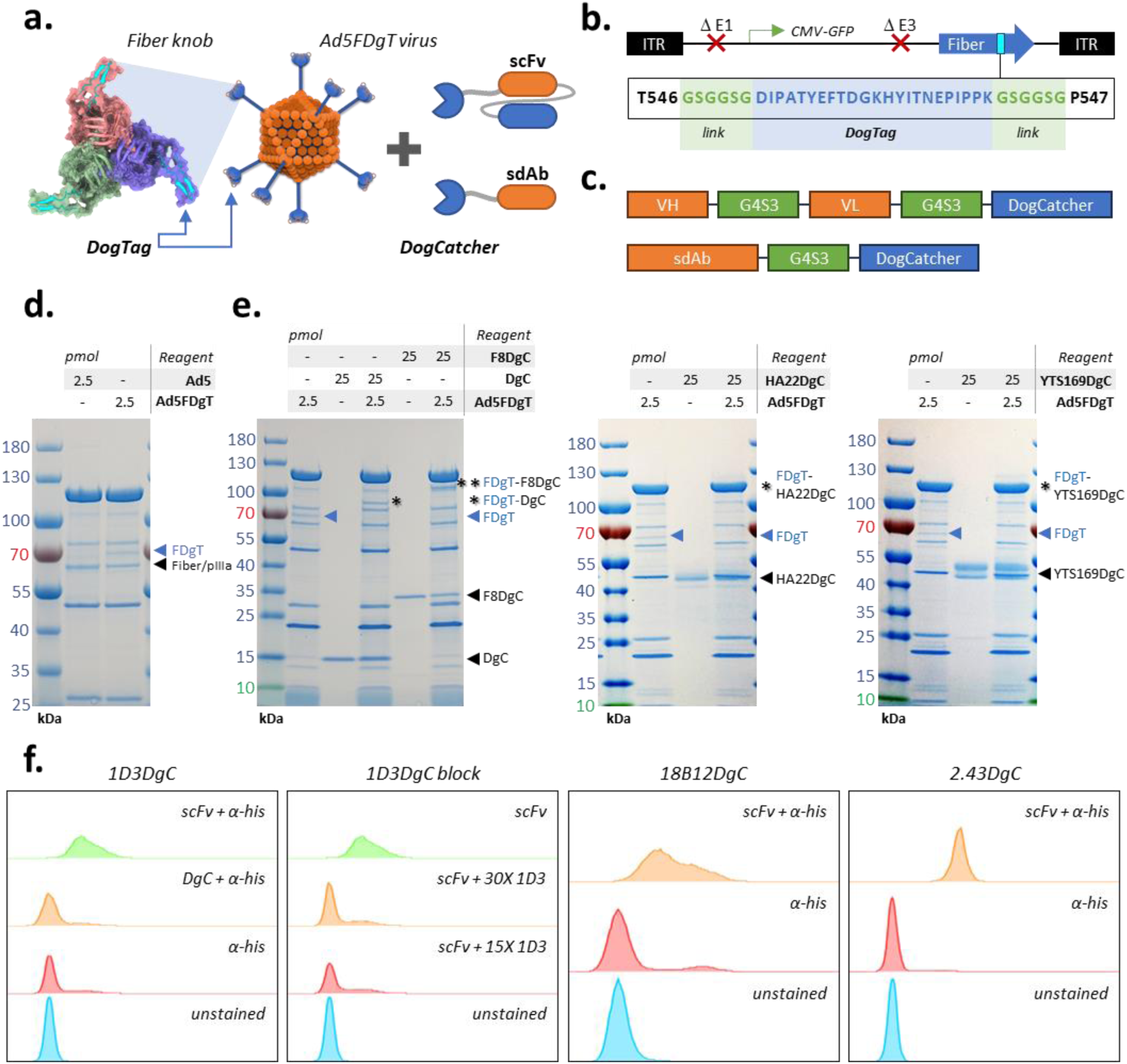
Development and characterization of Ad-Ab targeting. **a.** Schematic overview of system design. DogTag is genetically inserted into the Ad fiber knob (Ad5FDgT), while DogCatcher is fused to antibody species. Mixing of these reagents results in permanent linkage of the virus and antibody at the fiber knob locale. Fiber model generated with AlphaFold2 and visualized with UCSF ChimeraX^55,56^. **b.** Ad5FDgT genome overview. Ad5FDgT is based on an E1/E3 deleted Ad5 with the CMV promoter driving eGFP expression from the E1 region. DogTag is inserted with minimal flex linkers at the HI loop of the fiber knob domain. **c.** Antibody-DogCatcher fusion designs. Antibody domains are separated from DogCatcher by a flexible (G4S)_3_ linker **d.** SDS-PAGE analysis of control Ad5 and Ad5FDgT. In all cases the molar amount of antibody and virus refers to DogCatcher and DogTag, respectively. **e**. Gel shift analysis of Ad5FDgT binding to DogCatcher and antibody-DogCatcher fusions. Left, binding to bacterially produced DogCatcher and an sdAb-DogCatcher fusion. Middle and right, binding to representative scFv-DogCatcher fusions. **f.** Flow cytometry analysis of representative scFvs in murine splenocytes. 1D3DgC and 18B12DgC stains were gated on B220+ populations, while 2.43DgC staining was gated on the CD8+ population. 1D3DgC specificity was assessed by blocking with excess full length 1D3 antibody.

To pair with this vector, we developed several antibodies as DogCatcher fusions. We initially developed the aforementioned anti-mCD40 F8 sdAb as a DogCatcher fusion (F8DgC), but also developed single-chain fragment variable (scFv) fusions of commercial antibodies targeting murine and human B cell markers, including CD19 and CD20 (**Fig. 1c**). As controls and to assess targeting in an alternate cell type, we also developed scFv fusions targeting the murine T cell marker CD8α. To assess if these reagents were able to conjugate the virus fiber capsid protein, we co-incubated Ad5FDgT and different antibody fusions at room-temperature for approximately 2 hours, then ran SDS-PAGE gels to assess the degree of shifting in the fiber protein. Similar to a previous study, we found that the DogTag-DogCatcher pair was highly reactive and that all or nearly all of the fiber-DogTag protein reacted with the antibody fusions (**Fig. 1e**)^25^. Furthermore, we found these antibodies retained their ability to bind to the appropriate cell type, as determined by flow cytometry (**Fig. 1f**).

### In vitro and in vivo analysis of Adenovirus-antibody conjugates

To assess the ability of our Adenovirus-antibody (Ad-Ab) conjugates to achieve targeted gene transfer, we infected primary murine B and T cells with Ad-Ab decorated with antibodies targeting B or T cell restricted cell markers (CD40, CD19, CD20 and CD8α). As expected, we found that gene transfer enhancement occurred only when the appropriate antibody was conjugated on the virus – infectivity did not change when B cells were treated with a virus targeting CD8α, while T cell infection rates were not impacted by conjugation of the virus with CD40, CD19, or CD20 targeting antibodies (**Fig. 2b**). We also assessed this effect in primary human B cells, and similarly found that gene transfer enhancements only occurred in the presence of the appropriate targeting agent. In all cases we tested a variety of molar ratios of antibody-DogCatcher to virus-DogTag, and often observed a bell curve where very high and very low ratios of antibody resulted in lower gene transfer enhancements. This is consistent with the idea that too much antibody fragment may act as a competitive inhibitor, but too little antibody may not saturate the binding sites on the virus. We did observe several notable exceptions to this bell curve rule, especially in the human B cell samples where the highest ratio of antibody-DogCatcher to virus-DogTag resulted in the highest gene transfer enhancement. This may stem from differences in conjugation rates – some antibodies might require higher excesses to achieve full saturation of the virus surface. It is also possible that differences in receptor density on different cell types could play a role. Further work is needed to fully elucidate this effect. Our results were particular striking in murine T cells and human B cells, where Ad5FDgT without enhancing antibody was almost completely incapable of gene transfer.

**Figure 2:**
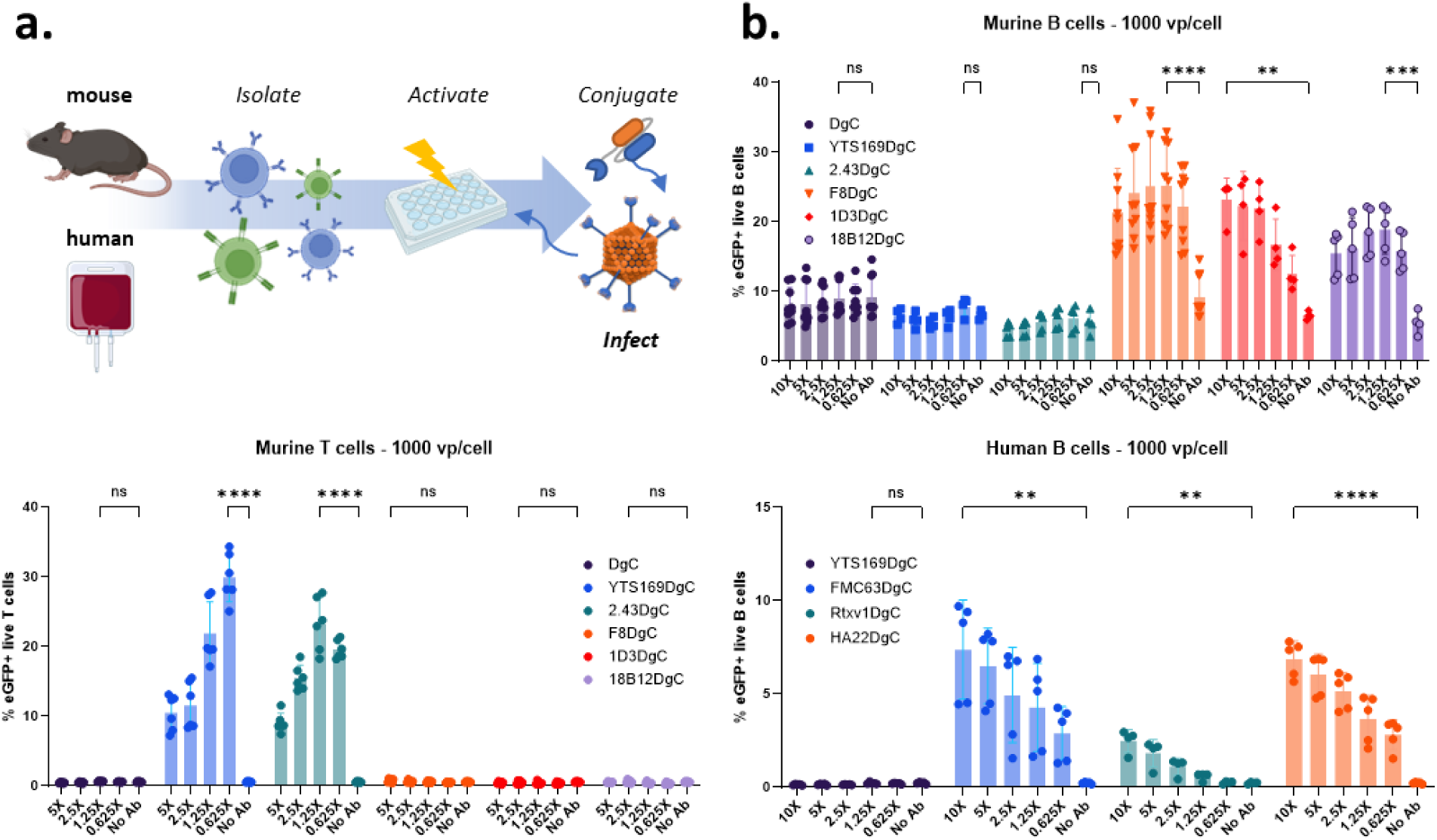
In vitro characterization of Ad-Ab targeting. **a.** Conceptual workflow. Primary lymphocytes are magnetically isolated from mouse splenocytes or human PBMCs then cultured with activating agents. On the day of infection Ad5FDgT is conjugated with the appropriate antibody at the indicated DogCatcher:DogTag molar ratios, then used to infect cells. **b.** Ad-Ab infectivity enhancement in primary murine B cells (top right), murine T cells (bottom left), and human B cells (bottom right). For each antibody, the group with the maximum mean infectivity was compared to the no antibody group using a standard unpaired two-tailed t-test. Welch’s correction was used in cases where the F-test revealed significant differences in variances. **DgC = DogCatcher**, n=9 replicates from 3 experiments in mouse B cells, n=6 from 2 experiments in mouse T cells. **YTS169DgC** = α-mCD8α scFv, n=4-5 from 2 experiments in mouse B cells, n=6 from 2 experiments in mouse T cells, n=5 from 3 experiments in human B cells. **2.43** = α-mCD8α scFv, n=4-5 from 2 experiments in mouse B cells, n=6 from 2 experiments in mouse T cells. **F8DgC** = α-mCD40 sdAb, n=9 from 3 experiments in mouse B cells, n=6 from 2 experiments in mouse T cells. **1D3DgC** = α-mCD19 scFv, n=4 from 2 experiments in mouse B cells, n=6 from 2 experiments in mouse T cells. **18B12DgC** = α-mCD20 scFv, n=4-5 from 2 experiments in mouse B cells, n=6 from 2 experiments in mouse T cells. **FMC63DgC** = α-hCD19 scFv, n=5 from 3 experiments in human B cells. **Rtxv1DgC (Rituximab)** = α-mCD20 scFv, n=4 from 2 experiments in human B cells. **HA22DgC** = α-hCD22 scFv, n=5 from 3 experiments in human B cells. Created with BioRender.com.

Encouraged by our *in vitro* results, we decided to determine if antibody conjugation could enhance specific gene transfer *in vivo*. We injected C57BL/6J mice retro-orbitally with Ad5FDgT alone or Ad5FDgT conjugated with 1D3DgC, an scFv targeting murine CD19, a B cell restricted cell marker (Ad5FDgT-1D3) (**Fig. 3a**). We scored splenic B and T cells for reporter gene expression three days later, and found a statistically significant enhancement of gene transfer in the B cell compartment, but not in the T cell compartment (**Fig. 3b**). This enhancement was found across several of the B cell subpopulations we scored, including memory B cells, marginal zone B cells, and follicular B cells (the gating strategy for these subpopulations can be found in **Fig. S1**). We did not observe enhancement in the plasmablast population, which may be due to lower CD19 expression on this cell type^29^. We also scored eGFP expression per milligram of tissue in several major organs, including the spleen, liver, lung, heart and kidneys. In the main site of Ad5 tropism *in vivo*, the liver, we observed a non-statistical trend towards decreased gene transfer in the Ad5FDgT-1D3 group, compared to the Ad5FDgT group (**Fig. 3c**). We also observed the Ad5FDgT-1D3 group showed statistically reduced lung and spleen eGFP expression, potentially indicating virus conjugation may reduce off-target gene expression in these tissues. Further work is required to elucidate the mechanism of these results.

**Figure 3:**
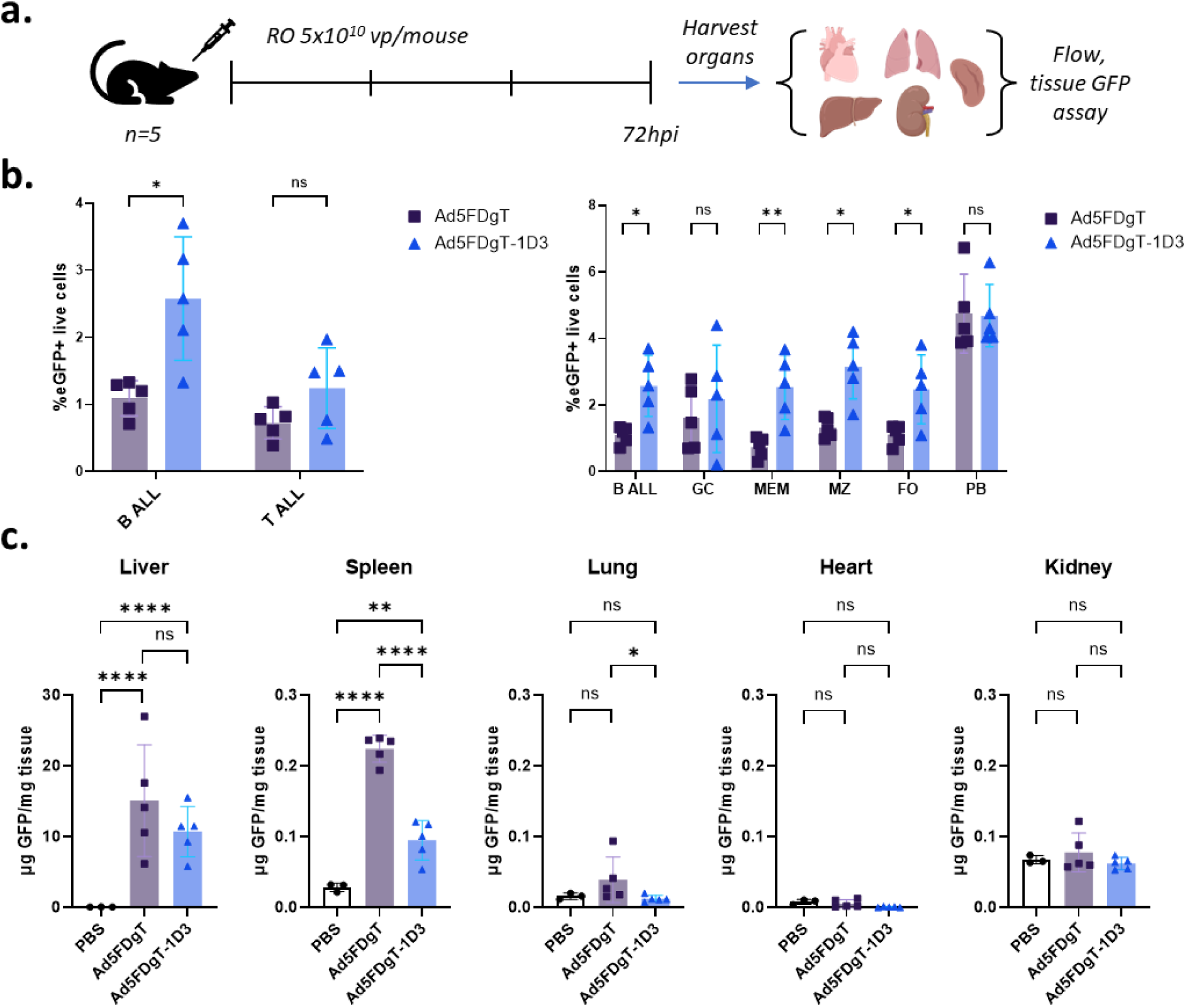
In vivo characterization of Ad-Ab targeting. **a.** Experiment design. C57BL/6J mice were injected RO with 5×10^10^ vp/mouse of either Ad5FDgT or Ad5FDgT conjugated with 1D3DgC. Major organs were harvested 72h later for eGFP expression analyses. **b.** Flow cytometry results. Left, eGFP expression in B and T cells. Right, eGFP expression in B cell subsets. GC = germinal center, MEM = memory, MZ = marginal zone, FO = follicular, PB = plasmablast. In all cases non-conjugated and conjugated groups were compared in each population using a standard unpaired two-tailed t-test. Welch’s correction was used in cases where the F-test revealed significant differences in variances. eGFP expression in mice injected with PBS was used to normalize the data in all cases. **c.** Quantitative eGFP analysis in major tissues. Tissues were homogenized in lysis buffer and eGFP expression was analyzed via fluorimetry. eGFP was normalized to total tissue protein content as assessed by BCA. In all cases ordinary one-way ANOVA with Tukey’s correction for multiple comparisons was used to compare groups where n=5 male mice. In the liver, lung and kidney values were log transformed to normalize variances. Created with BioRender.com.

### Development and analysis of purified single-component Ad-Ab complexes

Taken together, our data suggested that Ad-Ab complexes were a promising platform for controlling gene transfer and expression both *in vitro* and *in vivo*. However, for clinical translation and industrial production a single component vector without excess antibody fusion protein is desirable. This would also allow us to remove any potential confounding effects of excess antibody in our experiments. We therefore attempted to develop a workflow to purify and store purified Ad-Ab complexes (**Fig 4a**). At the lab scale, density gradient centrifugation is often used to prepare purified Ads. From cell lysates, we therefore used a single ultracentrifugation with cesium chloride gradients to purify Ad5FDgT. We then dialyzed this vector against 1X PBS to prevent any negative effects towards the virus conjugation from excess cesium chloride. We split this sample into equal thirds, and treated one fraction each with PBS, α-mCD19 1D3DgC, or α-mCD8α YTS169DgC. Conjugation was carried out at 25C for 1 hour, and vectors were ultracentrifuged a second time to separate free antibody from the virus conjugates. Importantly, observation of the vector bands did not reveal any obvious differences between virus treated with PBS and virus conjugated with scFvs (**Fig. 4b**). After spinning vector samples were dialyzed again against our standard 1X PBS with 10% glycerol and aliquoted and frozen at −80C.

**Figure 4:**
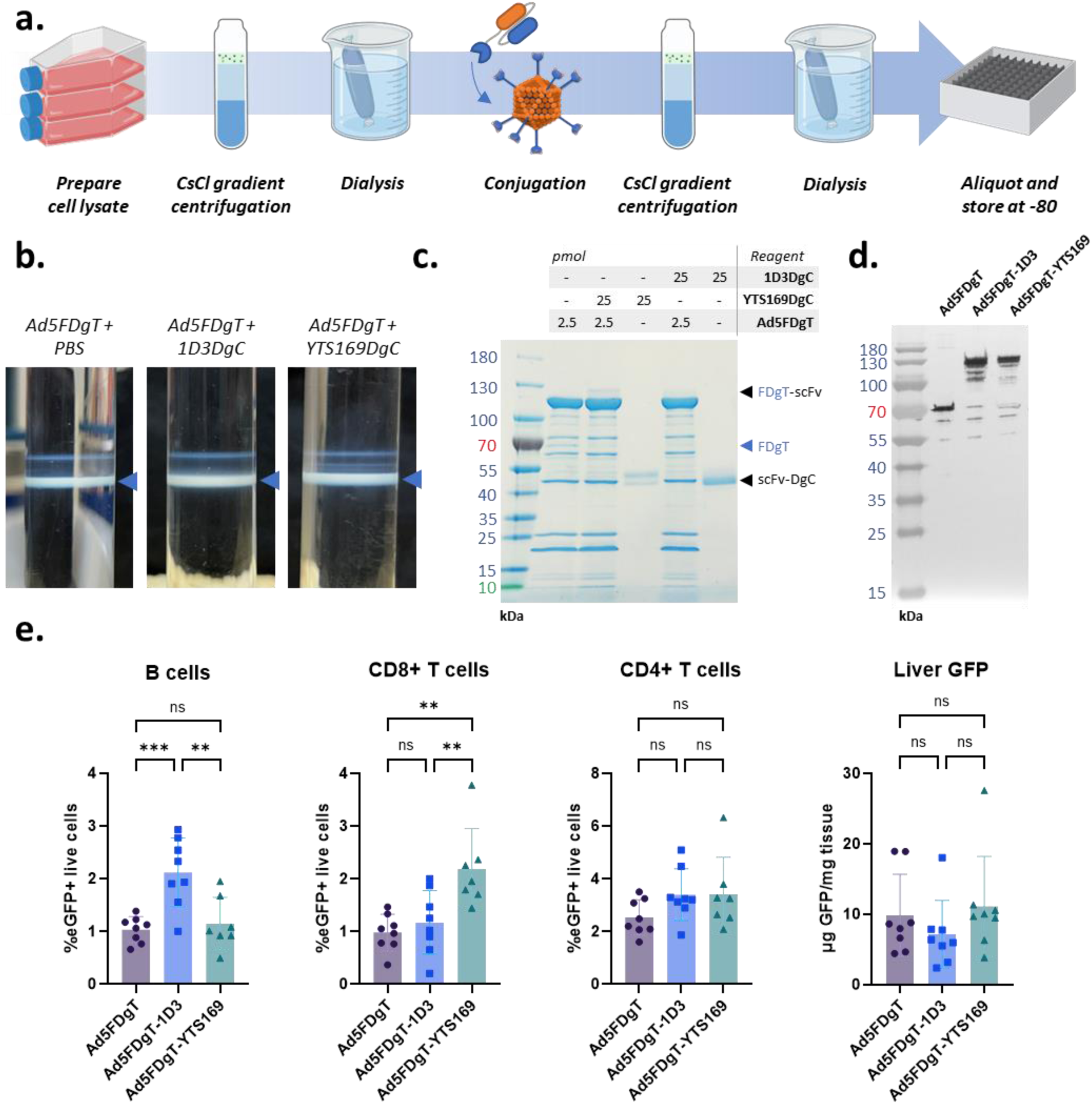
Development of purified Ad-Ab complexes. **a.** Purification workflow. Virus particles are purified from cell lysates via cesium chloride (CsCl) ultracentrifugation, then briefly dialyzed against 1X PBS to remove excess salts. Conjugation is then carried out, followed by a second CsCl purification, dialysis and final storage. **b.** Viral band images in CsCl gradients after second ultracentrifugation. **c.** SDS-PAGE analysis of purified Ab-Ab complexes. **d.** Western blot against the HAdV-C5 fiber tail of purified Ad-Ab complexes. **e.** In vivo analysis of purified Ad-Ab complexes. C57BL/6J mice aged 6-9 weeks were injected on three separate occasions with 5×10^10^ of the indicated vectors. In the first experiment n=2 female mice per group were used. In the second experiment n=2 male mice per group were used, and a single Ad5-FDgT-YTS169 sample was removed from analysis due to very low viability found during flow analysis. In the fourth experiment n=4 mice per group were used split equally between male and female mice. Three days later splenocytes were assessed for eGFP expression using flow cytometry. Livers were assessed for tissue eGFP expression as well. Data from all experiments were combined and analyzed using ordinary one-way ANOVA with Tukey’s correction for multiple comparisons. Created with BioRender.com.

We carried out several assays to determine the quality of these purified Ad-Ab complexes. Encouragingly, SDS-PAGE analysis revealed that the antibody-conjugated samples retained an identical protein band pattern to the PBS treated vector, with the exception of the expected complete shift in the fiber-DogTag band (**Fig. 4c**). Critically, we also were unable to observe any signs of free antibody. This result was further confirmed by a western blot analysis of the fiber protein, which revealed that the vast majority of fiber-DogTag reacted with the antibody-DogCatcher fusions (**Fig. 4d**). We then carried out *in vitro* analyses of these viruses and found that gene transfer enhancement from the purified and stored viruses was comparable to freshly prepared Ad-Ab complexes (**Fig. S2**).

Interestingly, we observed that different antibodies required different molar ratios of DogCatcher:DogTag to achieve optimal gene transfer *in vitro* (e.g., 1D3DgC was most potent in the 10X-2.5X range while YTS169DgC was most potent at 0.625X – see **Fig. 2**). We had hypothesized that this could be due to differences in the ability of each antibody fusion to link with the virus. However, with purified Ad5FDgT completely functionalized with YTS169DgC, we observed that gene transfer was lower than in our previous experiments with a 0.625X DogCatcher:DogTag (**Fig S2**). These results indicate that, at least for mCD8α targeting antibodies, an un-saturated capsid is superior to a saturated one. The Ad fiber presents as a trimer, with three copies of DogTag displayed in close proximity. Full functionalization of this trimer with antibody apparently results in lower gene transfer to murine T cells, potentially indicating excessive cross-linking of CD8α is sub-optimal for driving viral uptake. Additional work is needed to understand this effect and determine the minimum number of antibodies per capsid to achieve optimal targeting through CD8α. It would also be intriguing to determine if this effect is specific to T cells, or specific to CD8α.

We next endeavored to assess these purified complexes *in vivo* (**Fig. 4e**). As before, we injected naïve C57BL/6J mice retro-orbitally with either non-conjugated Ad5FDgT, Ad5FDgT-1D3, or Ad5FDgT-YTS169. We conducted three separate experiments, first in female mice, then male, and finally a mixed group. We again found that functionalization of the virus with 1D3DgC resulted in a roughly two-fold increase in gene expression to B cells compared to the parent vector (the gating strategy for the cell types described in this section can be found in **Fig. S3**). We found a similar enhancement in the CD8α+ T cell population, with YTS169DgC functionalization resulting in an approximately two-fold increase in gene expression. Critically, neither modification resulted in an increase in gene expression in off-target cell types – 1D3DgC functionalization did not increase gene expression in CD8α+ T cells, and YTS169DgC did not increase gene expression in B cells. We also did not observe statistical differences in groups separated by sex, confirming that our technology is applicable in both male and female mice (**Fig. S4**).

Interestingly, we did observe a slight, non-significant increase in gene expression in CD4+ T cells in mice injected with Ad5FDgT-1D3 and Ad5FDgT-YTS169 compared to Ad5FDgT. The potential mechanism of this result is not clear – activated CD4+ T cells do express an Fc receptor which could conceivably bind to the antibodies used to functionalize our vector, but our study uses scFvs lacking any Fc receptor binding^30^. Another explanation could be that incorporation of scFvs into the fiber increases the circulation time of the virus and allows slight increases in transduction of off-target cells. There may also be as-of-yet uncharacterized interactions between the various hematopoietic cells and scFvs or DogCatcher itself that are responsible. Additional work is required to understand this effect.

We also assessed liver eGFP expression per milligram of tissue as in our previous assay and selected two mice per group for tissue immunohistochemistry. We did again observe a weak, non-statistical decrease in liver gene expression in mice injected with Ad5FDgT-1D3, but this result was not true for mice injected with Ad5FDgT-YTS169. We also confirmed that there were not significant differences in liver eGFP expression in male and female mice (**Fig. S4**). We further confirmed these results using tissue immunohistochemistry and demonstrated that scFv vector functionalization did not result in any radical changes in overall gene expression across all organs (**Fig. S5**) – as described in numerous previous studies, the overall *in vivo* distribution of Ad5 to the liver is clearly dominated by binding of the liver to blood factors^31,32^.

## Discussion

Here we describe the development of a novel platform technology for targeting of Adenovirus to defined receptors using antibodies. We pair a simple, easy-to-produce vector with scFv fusions, enabling rapid design of vectors capable of efficient gene transfer into challenging cell types. The Ad-Ab system boosts gene transfer into cells moderately-to-completely resistant to Ad infection – in an exemplary test, murine CD8α+ T cells infected with unmodified Ad5FDgT at 25,000 vp/cell showed less eGFP expression than those infected with just 40 vp/cell of Ad5FDgT-YTS169 (data not shown). *In vitro*, Ad-Ab is thus a highly useful platform for gene delivery into primary cell types, especially in contexts where mixed cell populations are present or the cell type is very difficult to infect with conventional vectors.

Ad-Ab offers several key advantages over conventional Adenovirus targeting techniques. A classic example is the swapping of fiber knob domains from other Ad serotypes onto the Ad5 fiber tail and shaft, resulting in transfer of tropism from the swapped serotype. This strategy has been extensively used for hematopoietic stem cell (HSC) engineering, wherein swapping the Ad5 fiber for the fiber from Adenovirus serotype 35 results in vector targeting to CD46, which is present on HSCs^33–35^. Although this technique has been highly successful, there is a limited repertoire of serotypes available for fiber swapping, and many bind to proteins such as the coxsackie and adenovirus receptor, CD46, or desmoglein-2 which are expressed in many tissues^36^. Fiber swapping is thus a limited technique for targeting defined cell populations. In comparison, Ad-Ab can theoretically be targeted to any receptor for which an antibody has been described.

To overcome these limitations and target specific cell receptors, many groups have genetically engineered the Ad5 fiber to incorporate small peptides, ligands and sdAbs^18–20^. Although these approaches can be successful, extensive editing of the viral capsid often results in reduced titers and infectious particle ratios. In our own hands, we have generated numerous vectors genetically modified to express sdAbs at the fiber^19,37–39^. Although these efforts have yielded successes, we typically needed to attempt production of many sdAb variants to obtain a vector which was able to be grown to high titers. Additionally, there was no guarantee that successfully produced sdAb modified vectors would bind an accessible epitope *in vivo* – many vectors which displayed remarkable infectivity enhancements *in vitro* did not translate. The relative sparsity of available sdAbs also required us to develop novel variants for many antigen targets, a lengthy and costly process involving immunization of large animals. The ability to use the sequences of commercially available and characterized antibodies with validated *in vivo* effects to generate scFvs is thus a major advantage to the Ad-Ab platform.

To circumvent issues with viral titers and infectivity, many groups have attempted the use of adaptors to link antibodies or other targeting agents onto the vector surface – for example, our group has fused the soluble domain of the coxsackie and adenovirus receptor to various targeting agents in the past^40,41^. This protein is the endogenous receptor for Ad5 and binds the fiber protein with high affinity. A genetic fusion between coxsackie and adenovirus receptor and an antibody or ligand thus allows for vector retargeting. Another similar approach involves the use of a trimerized DARPin which binds the fiber knob fused to targeting DARPins^42^. While these systems show promise *in vitro*, the complex and challenging environment *in vivo* could break the interaction between the vector and the adaptor, leading to a loss of targeting and unexpected effects. Furthermore, such a two-component system may be challenging to translate clinically – a single component, permanently linked system which can be generated and processed in a similar manner to standard Ad vectors is thus a major advantage.

The use of molecular glue technology thus provides a solution to these issues and enables a simple targeting strategy. Other groups have also recognized this utility – SpyTag has been used to target lentivirus and was recently described for Ad targeting of a cancer cell line using an sdAb^43,44^. Our study validates and builds upon this work by swapping SpyTag for the highly reactive DogTag and demonstrating the use of scFvs for targeting primary cell types *in vitro* and *in vivo*, thus showing the potential of molecular glue driven targeting for clinical applications.

An inherent limitation of Ad5 based vectors, including our Ad-Ab system, is viral particle sequestration in the liver. This sequestration, at least for Ad5, has been demonstrated to be largely mediated by binding of Factor X to specific residues in the hexon protein^31,32^. Our group and others have thus developed numerous vectors capable of escaping liver sequestration through the use of targeted mutations in the hexon^45–47^. We anticipate the flexibility of the Ad-Ab system will allow us to combine the fiber targeting described here with additional capsid mutations – we have already been successful in developing a next-generation Ad-Ab vector with hexon modifications described by Atasheva et al to ablate liver and macrophage sequestration^47^. Similarly, a recent study incorporated DogTag in the hexon to link the virus with SARS-CoV-2 antigens to generate a novel vaccine platform^25^. Intriguingly, this group also reported that hexon functionalization with antigen blocked antibody mediated neutralization of the vector and binding of Factor X. Development of dual-tag modified vectors with DogTag at the hexon and fiber may also be of utility, as placing scFvs at both sites might lead to targeting through the fiber with simultaneous blocking of sequestration factors through the hexon.

Finally, we also previously demonstrated that SAd36 can be derivatized with SpyTag at various locales and used to deliver a SpyCatcher-Cas9 fusion protein to achieve gene editing^23^. Delivery of Cas9 as a protein rather than DNA may carry advantages such as reduced off-target effects due to its transient nature. Towards the combination of these systems, we have developed a variant of Ad5FDgT with SpyTag inserted at the C-terminus of the pIX protein, potentially allowing for targeting through fiber and delivery of Cas9 ribonucleoprotein through pIX. Further characterization of all the described vectors are of interest.

In total we present here a highly flexible and efficient platform for the transfer of genes to precisely targeted cell populations. We believe Ad-Ab adds to the toolbox of novel targeted vectors for gene delivery and may be of use for gene editing, especially *in vitro* in contexts where specificity and efficiency are paramount. With further modifications to the Ad-Ab capsid, we also anticipate strong utility for *in vivo* precision gene transfer. This system thus represents our initial steps on a path towards low-cost, targeted and safe gene therapy *in vivo* using Ad vectors.

## Methods

### Cell Lines

HEK293 (ATCC CRL-1573) cells were grown in Dulbecco’s Modified Eagle Medium/Ham’s F12 1:1 mixture supplemented with L-glutamine, 15mM HEPES, 10% fetal bovine serum (FBS) and 100U/mL penicillin-streptomycin. Cells were grown at 37C with 5% CO2 under sterile conditions.

### Viruses

A previously described first-generation E1/E3 deleted HAdV-C5 vector with the cytomegalovirus (CMV) promoter driving eGFP (Ad5.CMVeGFP) was used to generate Ad5FDgT^24^. The parent plasmid was cut with BarI (Sibenzyme) and BstBI (New England Biolabs) to release the fiber protein. Two PCR fragments were generated encoding the regions upstream and downstream of the HI loop domain with overlaps for the BarI and BstBI cut sites. A synthetic fragment encoding DogTag with short linkers and overlaps for the PCR fragment was synthesized by Integrated DNA Technologies (IDT). These three fragments were assembled with the cut Ad5.CMVeGFP backbone using NEB HiFi DNA Assembly (New England Biolabs), generating Ad5FDgT. This backbone was linearized using PacI and transfected into HEK293 cells for upscale. Viruses were purified and analyzed for viral particle concentration as previously reported^48^.

### Protein Constructs

The single domain antibody JPP-F8 was identified from the blood lymphocytes of two alpacas immunized with murine CD40 protein employing general methods previously reported in detail^49^. The JPP-F8 VHH was purified and evaluated by standard dilution ELISA (cite PMID: 33774040) and shown to bind murine CD40 with a sub-nM EC ^49^. pDEST14-F8DgC was derived by cloning of a synthesized DNA fragment (IDT) containing the camelid JPP-F8 single domain antibody followed by a (G4S)3 flexible linker into the SfoI site of pDEST14-DogCatcher (a gift from Mark Howarth, Addgene plasmid #171772; http://n2t.net/addgene:171772; RRID:Addgene_171772), and the resultant sequences encoded “6xHis-TEV-JPPF8-DogCatcher”^26^. pcDNA3.4-18B12DgC and pcDNA3.4-Rtxv1DgC were derived by cloning of synthesized DNA fragments (IDT) containing 18B12 and Rtxv1 scFvs together with a DgC fragment into the EcoRI and HindIII sites of pcDNA3.4-c-Fos scFv [N486/76] (a gift from James Trimmer, Addgene plasmid # 190560; http://n2t.net/addgene:190560; RRID:Addgene_190560), and the resultant sequences encoded “IL-2 signal sequence-18B12 scFV-(G4S)3-DgC-6xHis” and “IL-2 signal sequence-Rtxv1 scFV-(G4S)3-DgC-6xHis”^50^. pcDNA3.4-1D3DgC and pcDNA3.4-FMC163DgC were derived by cloning of synthesized DNA fragments containing 1D3 scFv and FMC163 scFv into the EcoRI and BamHI sites of pcDNA3.4-Rtxv1DgC, and the resultant sequences encoded “IL-2 signal sequence-1D3 scFv-(G4S)3-DgC-6xHis” and “IL-2 signal sequence-FMC63 scFv-(G4S)3-DgC-6xHis”. pcDNA3.4-HA22DgC, pcDNA3.4-1YTS169DgC pcDNA3.4-2.43DgC were derived by cloning of synthesized DNA fragments containing IGHV1-46 signal sequence followed by HA22 svFv, YTS169 scFv, and 2.43 scFv into XbaI and BamHI sites of pcDNA3.4-1D3DgC, and resultant sequences encoded “IGHV1-46 signal sequence-HA22 scFv-(G4S)3-DgC-6xHis”, “IGHV1-46 signal sequence-YTS169 scFv-(G4S)3-DgC-6xHis”, and “IGHV1-46 signal sequence-2.43 scFv-(G4S)3-DgC-6xHis”.

### Recombinant Protein Production

The plasmids pDEST14-F8DgC and pDEST14-DogCatcher were introduced into protein expression BL21(DE3)-RIPL E. coli cells. Single colonies were used to inoculate 25 mL starter LB containing 100 μg/mL carbenicillin and 50 μg/ml chloramphenicol grown at 37 °C overnight. The starter cultures were added to 500 ml fresh media without antibiotics, and cultures were grown at 37° C with shaking at 250 rpm for 2.5 hours. Protein expression was induced with 1 mM IPTG, and the cultures were incubated at 30 °C with shaking at 250 rpm for 4 hours. Cultures were centrifuged, and cell pellets were resuspended in lysis buffer (0.5 mM Tris, 0.3 M NaCl, 10 mM imidazole, 0.2% Triton X-100, 1 mg/ml lysozyme, 20 units/ml DNase I, 1 mM PMSF, and one complete mini EDTA-free protease inhibitor cocktail tablet per 10 ml) and incubated at 37° C for 30 minutes. The cell lysates were clarified by centrifugation at 32,000 rcf at 4° C for 30 minutes.

For mammalian recombinant protein production, 80-90% confluent 293T cells were transfected with pcDNA3.4-based protein expression plasmids in the presence of transporter 5 reagent (Polysciences, Inc.). The transfected cells were cultivated in DMEM medium containing 10% fetal bovine serum for 4 to 6 hours and switched to FreeStyle 293 medium for additional 4 to 5 days. The culture supernatants were collected and concentrated with Amicon Ultra-15 with 10K NMWL. The 6xHis-tagged recombinant proteins produced in bacterial and mammalian systems were purified using a HisPur Ni-NTA column with 20 to 40 mM imidazole washing buffer and 300 mM imidazole elution buffer, and eluted proteins were dialyzed in 10% glycerol in PBS with three buffer changes using 3.5KDa molecular weight cut-off Slide-A-Lyzer Dialysis Cassettes. Protein concentration was measured using BCA according to the manufacturer’s instructions (Thermo Scientific).

### SDS-PAGE and Western Blot

Whole virus protein analysis and conjugation analysis was carried out using SDS-PAGE. Viruses and proteins were incubated at a 2:1 ratio with 3x SDS sample buffer containing 187.5 mM Tris-HCl, 6% SDS, 30% glycerol, 0.125M dithiothreitol and 0.03% bromophenol blue at pH 6.8 for 15 minutes at 100° C. Samples were then loaded on a 4-15% gradient gel and resolved using a Criterion electrophoresis system (Bio-Rad). Staining was carried out using GelCode Blue according to the manufacturer’s protocol (Thermo Scientific).

Western blot analysis was carried out to detect the Ad5FDgT fiber protein and its modified/conjugated derivatives. The samples containing about 6 × 10^11^ viral particles were mixed 1:1 with the 2X Laemmli SDS-PAGE loading buffer containing 2-mercaptoethanol (Sigma) and heated in boiling water for 5 minutes to denature the viral proteins. The denatured samples were run on 4 - 20% Tris-Glycine Mini protein gel (Invitrogen) using Novex Tris-Glycine SDS Running Buffer (Invitrogen) as recommended by the manufacturer. The iBlot 2 Dry Blotting System (Invitrogen) was used to transfer electrophoretically resolved viral proteins from the gel to PVDF membrane (Invitrogen) as recommended by the manufacturer. We employed the iBind Western System (Invitrogen) for immunodetection of Ad5FDgT fiber proteins using the primary mouse monoclonal antibody 4D2 against the N-terminal fiber tail domain. 4D2 was a kind gift from Jeff Engler and was produced by the Hybridoma and Monoclonal Immunoreagent Core at the University of Alabama, Birmingham^51^. Secondary detection was carried out using anti-mouse IgG conjugated with Alkaline Phosphatase (AP) (Sigma). The protein bands bound with both primary and secondary antibody were developed with colorimetric AP substrate reagent kit (Bio-Rad) as recommended by the manufacturer.

### Primary Cell Culture

Primary mouse and human B cell culture and infection was carried out as previously described by our group^24^. Briefly, mouse B cells were magnetically isolated from splenocytes and cultured for 30h in RPMI 1640 supplemented with 10% FBS, 1X Nonessential Amino Acids, 1X sodium pyruvate and 1X 2-mercapto-ethanol. 50 µg/mL LPS was used as an activation agent. Infections were carried out overnight with 5×10^5^ cells in 50 µL LPS-free media with 5% FBS. Infected cells were then returned to 500 µL total volume with complete media and incubated for a total of 48 hours after infection prior to flow cytometry analysis.

Human B cells were magnetically isolated from healthy donor peripheral blood mononuclear cells and cultured for approximately 72 hours prior to infection according to previously described protocols^24,52–54^. For infections 2.5×10^5^ cells were infected in a total volume of 40 µL STEMMacs HSC Expansion media (Stem Cell Technologies) with 0.2% FBS for 3 hours, then transferred to 1mL complete media. Flow cytometry analysis was carried out 48 hours after infection.

Mouse T cells were isolated from splenocytes using the Miltenyi Pan T Cell Isolation Kit II according to the manufacturer’s instructions. Cells were activated for 1 hour prior to infection using the Miltenyi T Cell Activation/Expansion kit according to the manufacturer’s instructions. 1-2×10^6^ T cells were incubated with 200 µL anti-CD3/CD28 microbeads in 10mL RPMI 1640 supplemented as above and with 10ng/mL mouse IL-4. For infections 5×10^5^ cells were resuspended in 100 µL culture media and infected for 16-20 hours. An additional 150 µL complete media was then added and cells were analyzed via flow cytometry 48 hours after infection.

### Virus Conjugation

For *in vitro* experiments, virus was prepared at 2X the concentration required for infection and mixed with an equal volume of the required amount of antibody fusion protein diluted in 1X PBS. This mixture was incubated at room temperature or 25° C for about 2 hours prior to infection.

For *in vivo* experiments, the required amount of virus for all injections was incubated with the required amount of antibody without further dilution for 2 hours at room temperature or 25° C. 1X PBS was then added to bring the conjugated virus to the appropriate final volume.

### Flow Cytometry

For analysis of antibody-DogCatcher fusions, splenocyctes isolated from C57Bl/6J mice (about 1 million cells) were stained in 100 µL FACS buffer (1X PBS containing 0.5% BSA) with 0.5 µg of DgC, 1D3DgC, 18B12DgC, 2.43DgC, or PBS control at 4° C for 30 minutes. Samples were then washed with 2 ml FACS buffer and resuspended in 100 µl FACS buffer containing 0.5 µl FITC anti-His Tag antibody (BioLegend) and 0.5 µl Alexa Fluor® 594 anti-mouse B220 antibody (BioLegend) or 0.5 µl Alexa Fluor® 594 anti-mouse CD8a antibody (BioLegend). Samples were incubated at 4° C for 30 minutes, washed again in 2 ml FACS buffer then resuspended in 100 µl FACS buffer. For blocking studies, splenocytes were pretreated with 7.5 µg or 15.0 µg full length 1D3 antibody (BioLegend) at 4° C for 30 minutes before the addition of 1D3DgC.

For analysis of *in vitro* infected murine B cells, samples were stained with anti-CD19 (Invitrogen #RM7717) and Fixable Far Red live/dead dye (Invitrogen). Samples were gated as singlets/live/CD19+. Human B cells were stained with anti-CD19 (BioLegend) and Fixable Far Red dye. Samples were gated as above. Mouse T cells were stained with anti-CD8α (BioLegend) and Fixable Far Red dye or Sytox Red dye (Invitrogen). Samples were gated as singlets/live/CD8α+.

For *in vivo* studies, spleens were processed into single cell suspensions and resuspended in 1mL 1X PBS with 2% FBS. 100uL of the suspension was used for analysis and was pre-incubated with 10 µL Fc Blocking Reagent (Miltenyi) for 10 minutes prior to staining. In all cases live/dead discrimination was carried out using Fixable Far Red dye. For the first *in vivo* assay (**Fig. 3**), B cells were gated as singlets/live/CD19+/CD3-, while T cells were gated as singlets/live/CD3+/CD19-. Germinal center B cells were gated as GL7+/CD95+/IgDlo/CD38lo, marginal zone as IgM+/IgDlo, follicular as IgMlo/IgD+, memory as IgDlo/GL7-/CD38+, and plasmablasts as CD138+/IgD-. All B cell subsets were first passed through a singlets/live/CD19+ gate prior to further analysis.

For the second *in vivo* assay (**Fig. 4**), B cells were gated as singlets/live/CD19+/CD3-/CD22+/CD4-/CD8-. CD8α+ T cells were gated as singlets/live/CD8α+/CD19-/CD3+/CD22-/CD4-. CD4+ T cells were gated as singlets/live/CD4+/CD19-/CD3+/CD22-/CD8α-.

Mouse antibodies used were as follows: CD19 (Invitrogen or Miltenyi), CD8α (BioLegend), CD3 (Invitrogen), GL7 (BioLegend), CD95 (Invitrogen) IgD (Invitrogen), CD38 (Miltenyi), IgM (Invitrogen), CD138 (Invitrogen), CD22 (BioLegend), CD4 (Invitrogen).

All analysis was carried out using FlowJo 10.10.0.

### Animal Studies

C57Bl/6J mice were acquired from the Jackson Laboratory and housed in a pathogen-free environment. Mice aged 6-9 weeks were injected retroorbitally with 5×10^10^ viral particles in 150 µL total volume and sacrificed roughly 72 hours later via anesthetization with Avertin followed by cervical dislocation. In the first *in vivo* study, one-half of each spleen was used for flow cytometry analysis, while the remaining half, liver, lungs, kidneys and heart were snap frozen in liquid nitrogen. Organs were thawed on ice and processed for eGFP quantification using a Fluorimetric GFP Quantitation Kit (Cell Biolabs) per the manufacturer’s instructions. Tissues were homogenized in the included lysis buffer supplemented with 10% Proteinase Inhibitor Cocktail (Sigma) using an Omniprep 96 automated homogenizer. Tissue lysated were then centrifuged at 4000-5000 rpm to remove cell debris and supernatants were transferred to clean tubes. Supernatants were diluted as appropriate then aliquoted into 96-well plates for eGFP quantification. Total solution eGFP was normalized against total solution protein from a BCA assay of the lysates performed according to the manufacturer’s instructions (Thermo Scientific).

In the second *in vivo* study, the whole spleen of each animal was used for flow cytometry and whole livers were used for tissue eGFP quantitation as above. All experiments were approved by the Institutional Animal Care and Use Committee of the Washington University in St. Louis School of Medicine (Protocol #22-0360) and were performed in accordance with the National Institutes of Health Guide for Care and Use of Laboratory Animals. All efforts were made to minimize suffering and the total number of animals used in the study.

### Purification of Ad-Ab complexes

Cells from 20X infected T175 flasks were harvested after the development of cytopathic effect and centrifuged at 1200 rpm for 5 minutes. Supernatants were removed and cell pellets were frozen at - 80° C. Samples were subjected to 3 freeze-thaw cycles in room-temperature water and dry ice to lyse the cells, then centrifuged at 4000 rpm for 10 minutes to clarify the lysates. Supernatants were then ultracentrifuged on cesium chloride gradients for 2 hours at 25,000 rpm at 4° C. Viral bands were harvested and dialyzed twice against 1X PBS. Virus was removed from dialysis and viral particles were quantified as above. Virus was then split into three equal aliquots and treated with either PBS, two-fold molar excess of 1D3DgC, or two-fold molar excess of YTS169DgC for 1 hour at 25° C. Conjugated viruses were ultracentrifuged again on cesium chloride gradients for 1.5 hours at 25,000 rpm at 4° C. Viral bands were harvested and dialyzed three times against 1X PBS with 10% glycerol, then frozen at −80° C.

### Statistical Analyses

Specific methods used for analysis are noted in the corresponding Figure legends. In all cases statistical analysis was carried out using GraphPad Prism 10. A p value of < 0.05 was used and significance is indicated as *: p < 0.0322, **: p < 0.0021, ***: p < 0.0002, **** p < 0.0001, ns: not significant. In all cases error bars correspond to standard deviation.

## Data availability statement

Plasmids used in this study are to be deposited in AddGene. Flow cytometry data is available upon reasonable request. All other data can be found in the main text or Supplemental.

## Acknowledgements

The authors would like to thank Mark Selby, Rosa Romano, and Hongil Park of Walking Fish Therapeutics, Inc for their contributions to human B cell culturing and analysis. This work was supported by National Institute of Health grants 1R21EB033459-01A1 and 1R01AI174270-01A1 awarded to David T. Curiel, 1R21HL166887-01A1 awarded to Zhi Hong Lu, and T32HL007088-45 awarded to Stephen Oh. Additional funding for this work was provided by Walking Fish Therapeutics, Inc through award P21-04949 to Zhi Hong Lu.

## CrediT author contributions

**Paul J. Rice-Boucher:** Conceptualization, formal analysis, investigation, visualization, methodology, writing – original draft. **Elena A. Kashentseva:** Methodology, investigation. **Igor P. Dmitriev:** Methodology. **Hongjie Guo:** Methodology, investigation. **Jacqueline M. Tremblay:** Resources. **Charles B. Shoemaker:** Resources. **David T. Curiel:** Conceptualization, supervision, writing – review and editing. **Zhi Hong Lu:** Supervision, conceptualization, formal analysis, visualization, investigation, methodology, writing – review and editing.

