## Supplemental Information for "In vivo targeted gene delivery using Adenovirus-antibody molecular glue conjugates"

### Supplementary

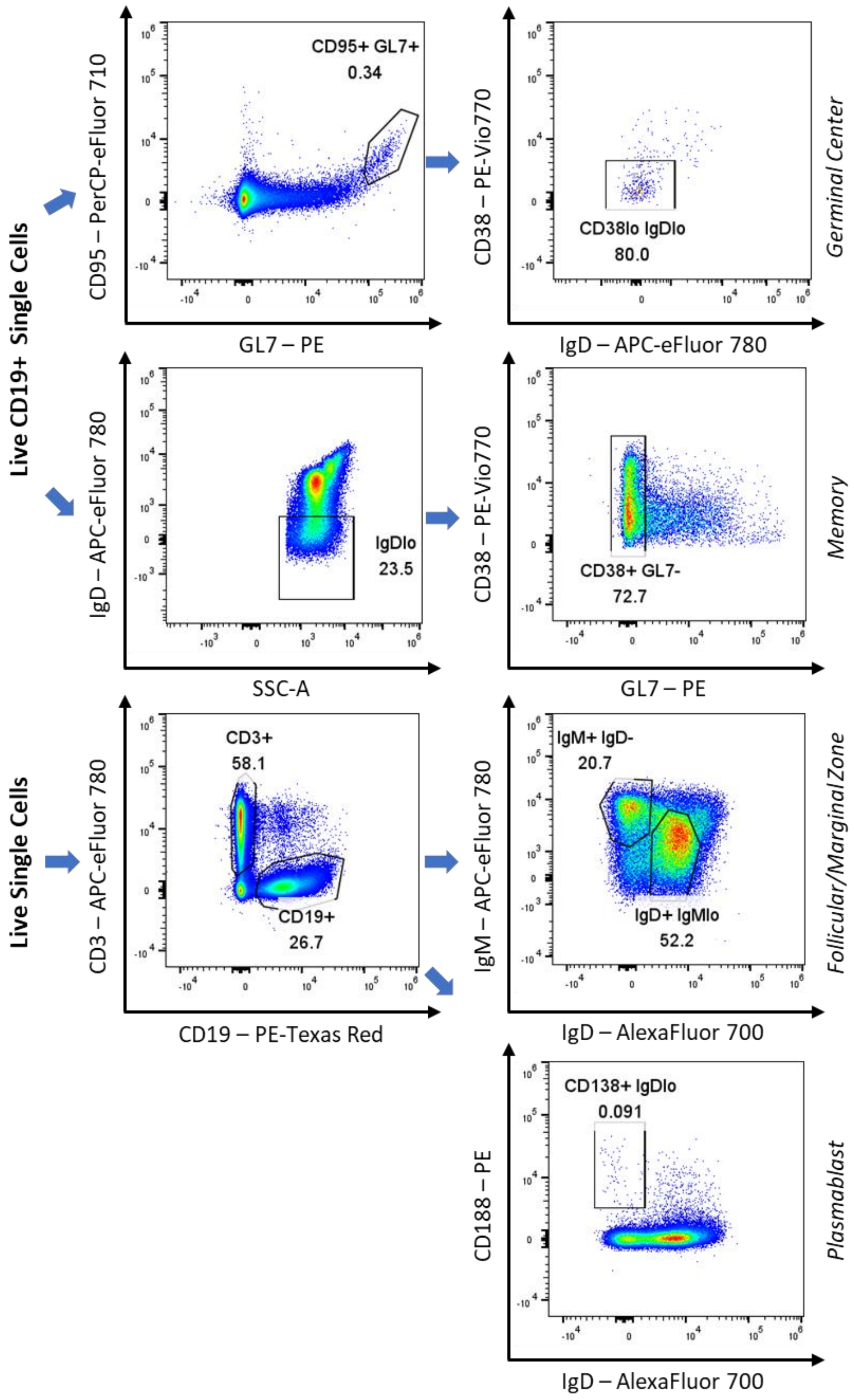


**Figure S1:** Flow cytometry gating strategy for cell populations defined in main Figure 3.

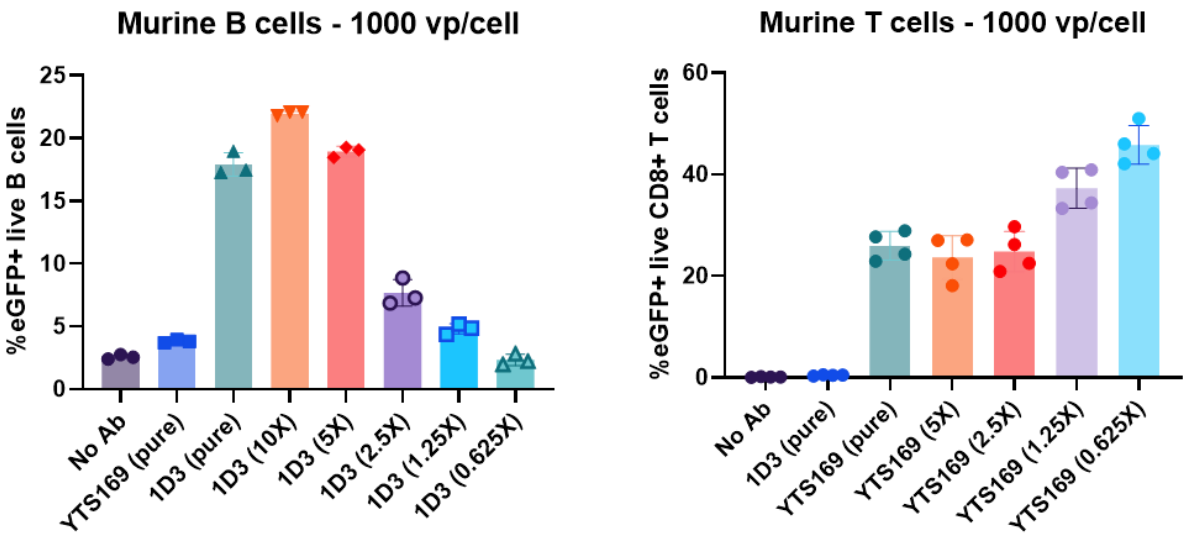


**Figure S2:** *In vitro* analysis of purified Ad-Ab complexes in murine B and T cells. Purified complexes are denoted as "scFv name (pure)" while freshly prepared complexes are denoted as "scFv name (molar ratio DgT:DgC). N=3-4 from 1 experiment.

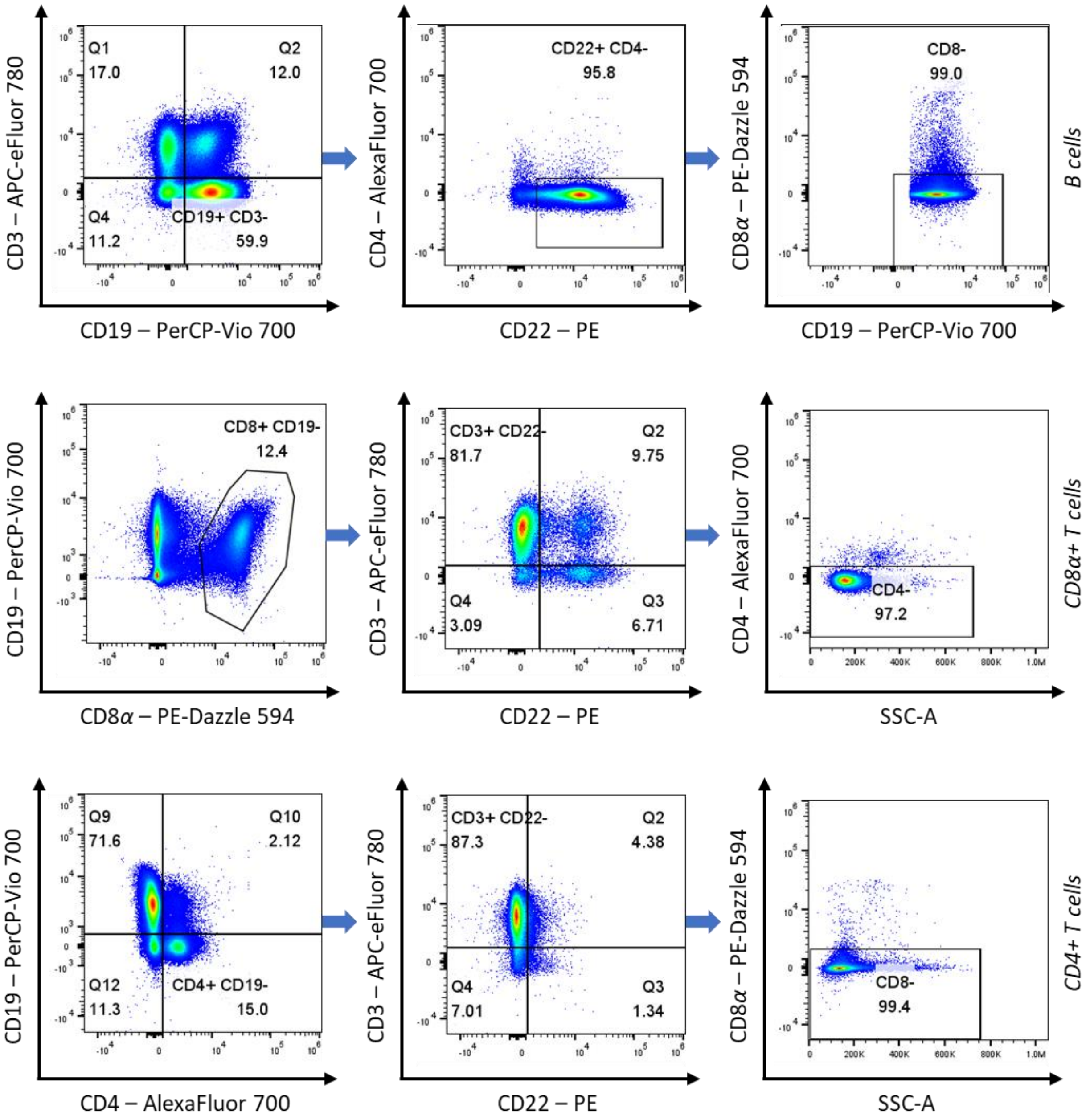


**Figure S3:** Flow cytometry gating strategy for cell populations described in main Figure 4. All populations were first gated on live single cells. We note that CD8α+/CD19- cells were gated on "curly quads" for analysis, which did not display correctly for the purposes of this graphic. We therefore substituted a polygon gate to visually demonstrate our gating strategy here.

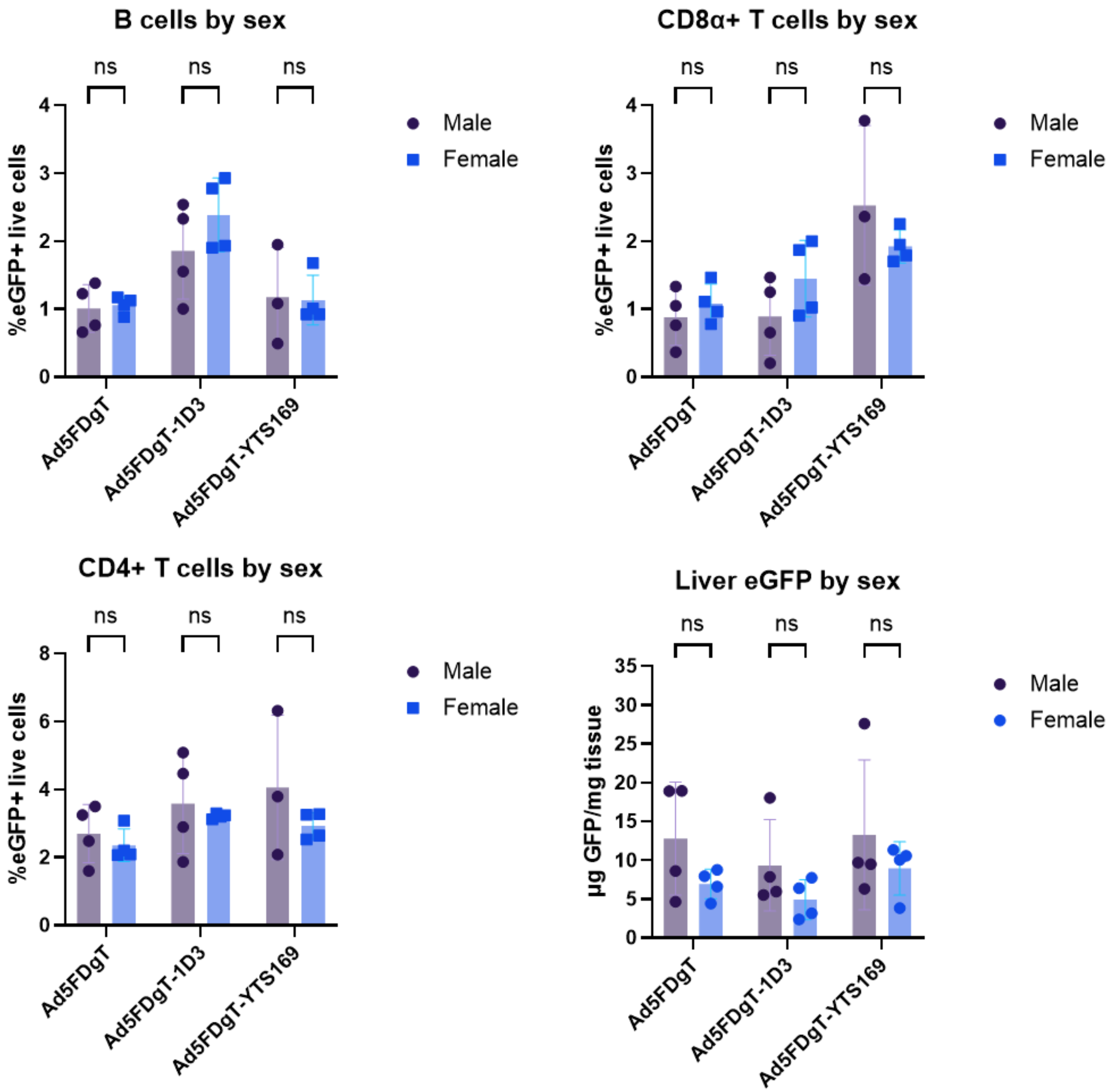


**Figure S4:** *In vivo* analysis of Ad-Ab targeting by sex. Data from Figure 4 was re-analyzed by sex using multiple unpaired t-tests with the Holm-Šídák correction for multiple comparisons. Sample sizes are described in Figure 4.

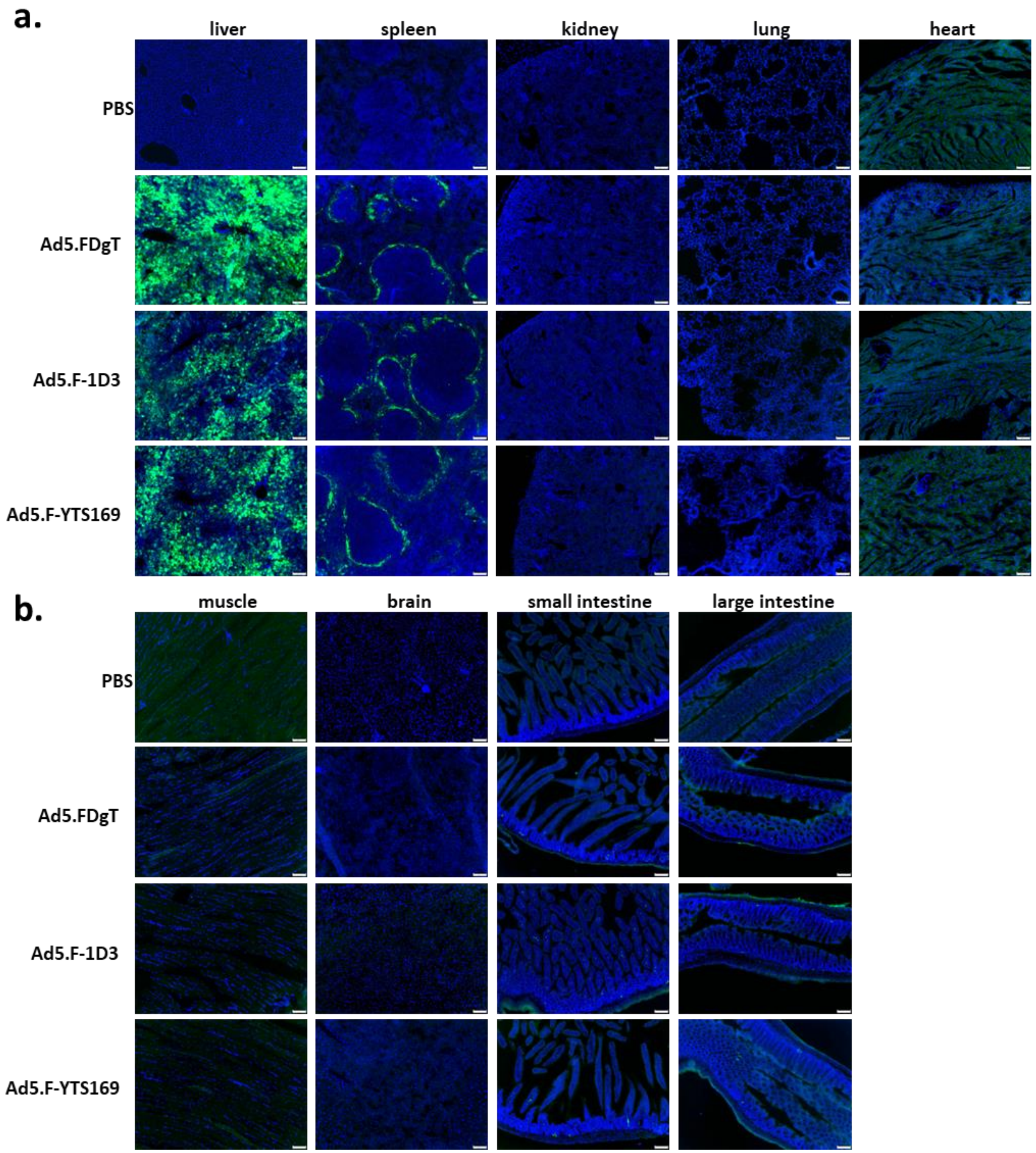


**Figure S5:** Nine organ tissue immunohistochemistry analysis. **a**. eGFP expression in liver, spleen, kidney, lung, and heart. **b**. eGFP expression in muscle, brain, small intestine, and large intestine. One male and one female mouse from each group described in **Fig. 3E** were perfused and organs harvested, then assessed for eGFP expression. Representative images from each group were picked and shown here.

**Supplemental Methods:**

**Tissue Immunohistochemistry:**

C57Bl/6J mice were obtained from Jackson Labs and injected with the indicated vectors as described in the main (one male and one female mouse per group). Three days after virus administration, mice were anesthetized with Avertin and perfused via the left ventricle with 10% neutral-buffered formalin. Lung was further inflated by injecting formalin solution into the trachea, closing the trachea by ligature, and then processing as below. Harvested organs were postfixed in formalin at room temperature for 2–4 h and cryopreserved in 30% sucrose in PBS at 4 °C overnight. Treated tissues were embedded in NEG50 (Thermo Scientific) and frozen on dry ice. All mouse tissues were cryosectioned at 16 µm. Frozen section slides were air-dried for 10 min, washed three times in PBS, blocked with protein block solution (5% donkey serum and 0.1% Triton X-100 in PBS) for 1 h, incubated at room temperature in protein block containing chicken anti-GFP 1:400 (ThermoFisher) for 1 hour, washed three times in PBS, incubated with corresponding  Alexa Fluor 488-conjugated secondary antibody 1:400 (Jackson ImmunoResearch Laboratories), and counterstained with SlowFade Gold Antifade mounting reagent with 4′,6-diamidino-2-phenylindole (DAPI) (Life Technologies). Immunofluorescence images were collected using an Olympus BX63 microscope equipped with an FVII digital camera (Olympus America).
